# Epstein-Barr Virus in HIV-Associated Conjunctival Squamous Cell Carcinoma and Transcriptional Reprogramming of Conjunctival Epithelial Cells

**DOI:** 10.64898/2026.10.20.760218

**Authors:** Hirotomo Dochi, Janardhan Avilala, Yutaro Yamada, Masayuki Takahira, Satoru Kondo, Tomokazu Yoshizaki, Maria Morales, Ramsy Abdelghani, David Becnel, Mark Sides, Gilbert F. Morris, Wenke Feng, Hong Liu, Zhen Lin

## Abstract

The involvement of Epstein-Barr virus (EBV) in conjunctival squamous cell carcinoma (cSCC) remains incompletely defined. This study integrated analyses of primary tumors with an experimental model of the conjunctival epithelium. Five Nigerian tumors from patients with HIV and three Japanese tumors from patients without HIV were examined by EBV-encoded RNA in situ hybridization (EBER-ISH), EBV DNA PCR, and virome analysis of RNA sequencing data. EBER localized to epithelial tumor cells in four Nigerian specimens and none of the Japanese specimens. EBV DNA and RNA were each detected in three Nigerian tumors, with every EBER-positive tumor supported by at least one additional assay. Human telomerase reverse transcriptase-immortalized conjunctival epithelial cells infected with Akata-GFP-EBV expressed latency-associated and lytic viral genes. Comparison of three infected and three control cultures identified 96 differentially expressed host genes and 23 enriched Hallmark gene sets at a false discovery rate below 0.05. RT-qPCR confirmed increased HES1, CXCL8, CXCL2, and LOX and decreased FGFBP1, CSTA, SAPCD2, and TMEM40 expression. Prominent responses involved inflammatory signaling, epithelial remodeling, metabolism, and cell-cycle regulation. These findings identify tumor-cell-associated EBV in a subset of HIV-associated cSCC and establish an epithelial model for investigating infection-associated host responses.

## INTRODUCTION

Conjunctival squamous cell carcinoma (cSCC) is an epithelial malignancy of the ocular surface with a marked association with human immunodeficiency virus (HIV)-related immune dysfunction in affected populations. A large South African cohort linked cSCC risk to immunosuppression and geographic proximity to the equator, highlighting the interaction between host immunity and environmental exposure ^1^. Identifying infectious contributors to this disease may help explain its etiologic diversity and reveal biological processes relevant to prevention and treatment.

Epstein-Barr virus (EBV) is a compelling candidate in this setting. Studies of ocular surface squamous neoplasia in Botswana and Zambia detected EBV in lesions from populations with a high prevalence of HIV infection ^2,3^. A subsequent case-control study in Zimbabwe identified an association between EBV DNA and cSCC ^4^. These observations motivate a complementary approach that localizes viral signals within tumor tissue and examines the response of conjunctival epithelial cells to EBV infection. Together, these approaches can connect clinical viral detection with experimentally accessible epithelial biology.

The epithelial context is particularly important. Organotypic studies have demonstrated EBV infection in stratified epithelium, while experimental work has connected the viral latent membrane protein LMP1 and host YAP/TAZ signaling to epithelial differentiation and viral-state regulation ^5–7^. An infection system derived from primary human conjunctival epithelium therefore offers an opportunity to characterize viral transcription and host responses in the cell type relevant to the ocular surface, rather than examining an isolated viral gene.

This study investigated EBV in cSCC specimens from Nigeria and Japan by integrating tumor-cell EBER-ISH with EBV DNA PCR and RNA-seq-based virome analysis. The experimental component established Akata-GFP-EBV infection in hTERT-immortalized human conjunctival epithelial cells and profiled both viral transcription and the host response. The study combines tissue-based evidence with a controlled epithelial cell model to identify candidate mechanisms linking EBV infection to inflammatory signaling and epithelial remodeling.

## Materials and Methods

### Clinical specimens

Eight existing formalin-fixed, paraffin-embedded (FFPE) cSCC specimens were analyzed. Five Nigerian specimens from patients with HIV were purchased from Audubon Bioscience, and three Japanese specimens from patients without HIV were provided by collaborating investigators from existing biobank collections. Diagnoses were confirmed by pathology review. Age, sex, geographic origin, and HIV status were recorded. Manuscript identifiers cSCC-1 – cSCC-5 denote Nigerian tumors and cSCC-6 – cSCC-8 denote Japanese tumors (Figure 1A).

**Figure 1.**
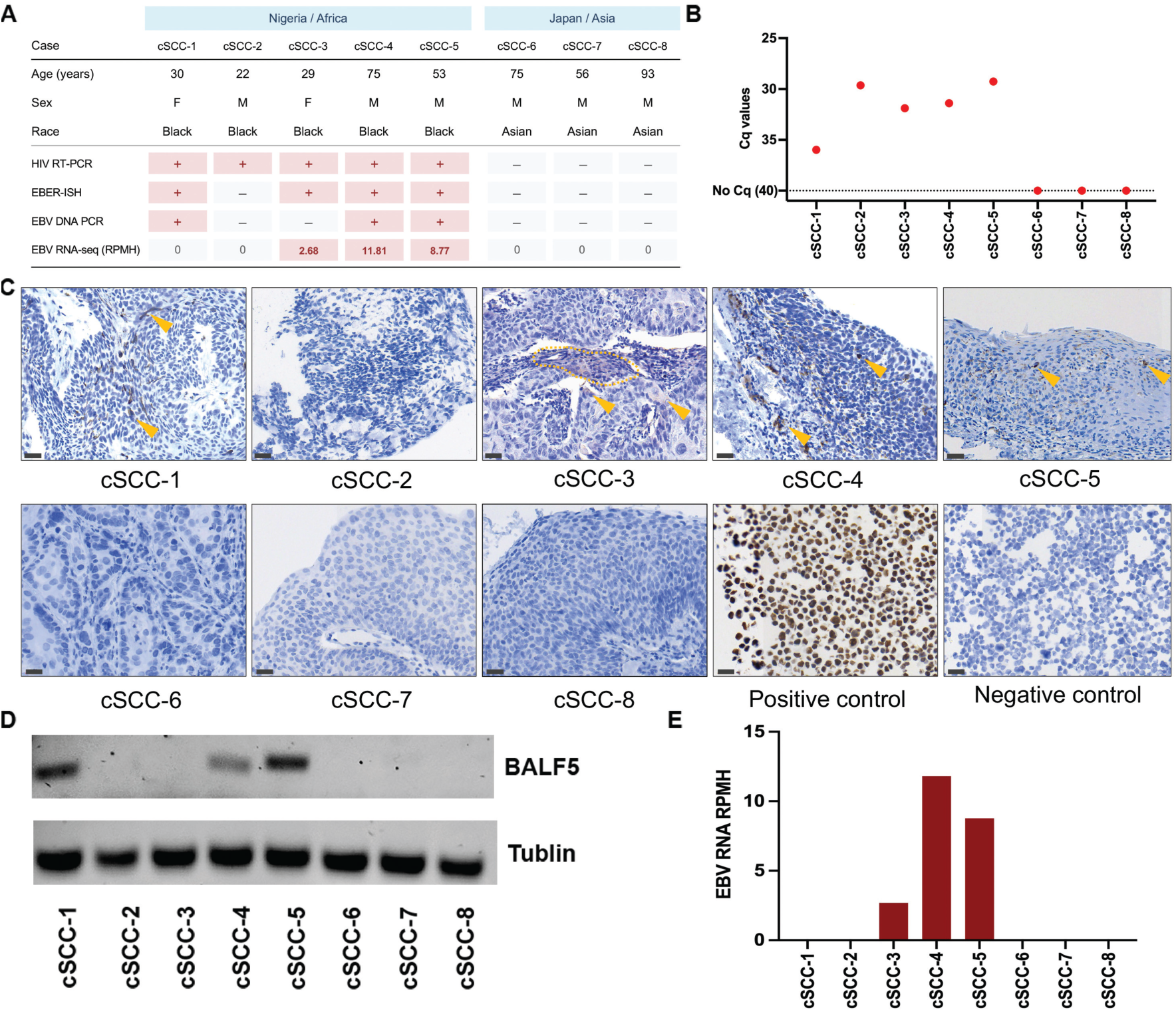
Tumor-cell EBV detection in primary conjunctival squamous cell carcinoma. (A) Clinical cohort and concordance matrix for virus-screening assays. The EBV RNA row reports reads per million mapped human reads (RPMH). (B) HIV RT-qPCR assay. (C) Representative EBER-ISH images showing nuclear signal in epithelial tumor cells, negative specimens, and assay controls. Scale bars, 30 μm. (D) EBV genomic DNA PCR targeting BALF5 and tubulin. (E) EBV RNA-seq abundance in the eight tumors, expressed as RPMH and shown on a linear scale. RPMH equals the number of qualifying species-aware EBV read ends divided by the mapped human read-end count and multiplied by 10⁶. Specimen identifiers are consistent across panels. cSCC-1 to cSCC-5 are Nigerian tumors, and cSCC-6 to cSCC-8 are Japanese tumors.

### Ethics approval and consent

All specimens were obtained from existing biobank collections with prior donor informed consent permitting research use. The study was approved by the Tulane University Institutional Review Board (Protocol# 2026-1466).

### EBER in situ hybridization

Five-micrometer FFPE sections were analyzed using the HistoSonda EBER XISH Probes kit (American MasterTech), with diaminobenzidine detection and hematoxylin counterstaining. EBV-positive and EBV-negative controls were included. Slides were scanned with a Motic EasyScan Pro 6 at 40× magnification. EBER positivity required discrete nuclear signal in morphologically identifiable epithelial tumor cells. Infiltrating lymphocytes were assessed separately. Tumor-cell EBER-ISH defined the primary EBV classification ^8^. Detailed staining procedures are provided in Supplementary Methods S1.

### Viral nucleic-acid assays

DNA and RNA were extracted using the Qiagen AllPrep kit. Clinical HIV status was evaluated additionally using the HIV TaqMan RT-PCR kit (Norgen Biotek, Cat# TM33750). EBV DNA was detected by SYBR Green qPCR targeting BALF5, with gel electrophoresis of amplicons and human genomic tubulin as an extraction/amplification control. HPV DNA was tested using Norgen multiplex endpoint PCR panels EP32200 and EP31500, collectively covering 18 genotypes but not HPV41 or HPV122. Positive, no-template, and internal human-DNA controls were included in HPV testing. Cycling conditions and HPV-panel coverage are detailed in Supplementary Methods S2. Primers are listed in Supplementary Table S1.

### Conjunctival epithelial cells and EBV infection

Primary human conjunctival epithelial cells (HConEpiC; ScienCell, Cat# 6630) were cultured in corneal epithelial medium at 37 °C and immortalized by lentiviral hTERT delivery with hygromycin selection. HConEpiC-hTERT cells received 100 µL cell-free Akata-GFP-EBV (6.7 × 10⁵ Raji IU/mL) in 2 mL medium at an MOI of 0.17. Mock-infected controls underwent matched handling without infectious virus. Three independently EBV-infected cultures and three independently maintained uninfected control cultures were harvested five days after infection or mock exposure. Infected cells were enriched by GFP-based fluorescence-activated cell sorting. Reactivation experiments used PMA (20 ng/mL) and sodium butyrate (3 mM) for 48 h. Additional details are in Supplementary Methods S3.

### RT-qPCR and immunoblotting

cDNA was prepared using iScript gDNA Clear cDNA Synthesis Kit and analyzed with iQ SYBR Green Supermix on a Bio-Rad CFX96. Melt curves assessed amplification specificity, and technical triplicates were averaged within biological samples. GAPDH was the RT-qPCR reference gene. Expression was calculated by the 2^−ΔΔCt^ method relative to uninfected controls. For undetectable EBV targets in controls, Ct = 40 was assigned to calculate the displayed estimates. This was an imputed, not measured, Ct. Figure 2B used post-sort infected cells without PMA or sodium butyrate. Immunoblots of whole-cell extracts assessed EBNA1, Zta (BZLF1), EAD/BMRF1, and VCA/p18, with GAPDH as the loading control and anti-IgG-induced Akata cells as the positive control. Signals were detected by chemiluminescence or infrared imaging. Only uniform image-wide or panel-wide adjustments were applied. Reaction, antibody, and imaging details are provided in Supplementary Methods S4-S5.

**Figure 2.**
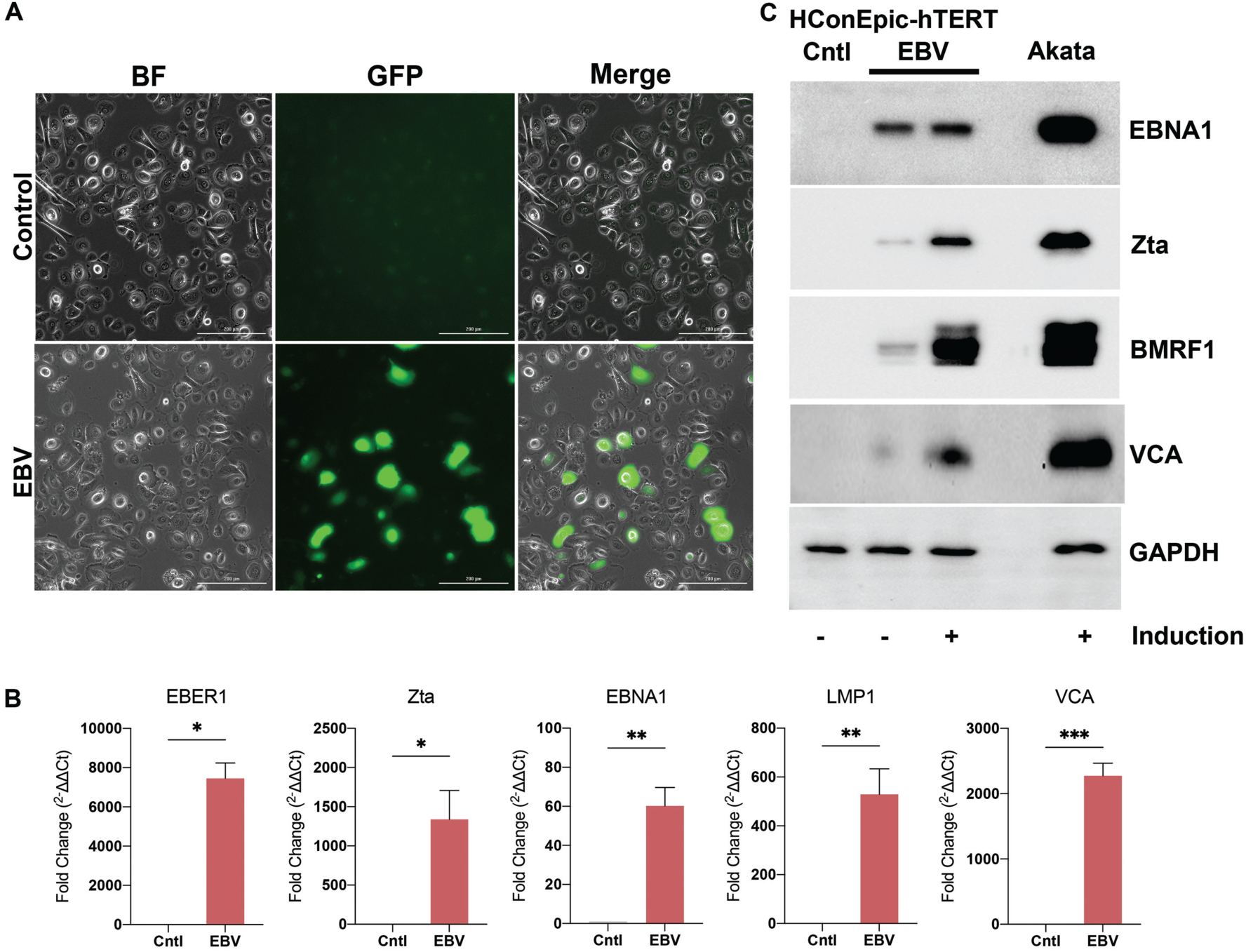
Establishment and validation of EBV infection in human conjunctival epithelial cells. (A) Bright-field, GFP fluorescence, and merged images of control and Akata-GFP-EBV-infected HConEpiC-hTERT cells before fluorescence-activated cell sorting (FACS). (B) RT-qPCR analysis of EBER1, BZLF1 (encoding Zta), EBNA1, LMP1, and VCA/BcLF1 in FACS-sorted GFP-positive EBV-infected cells and uninfected controls. The infected cells were not treated with PMA or sodium butyrate. GAPDH was the internal reference gene, and relative expression was calculated by the 2^−ΔΔCt method relative to uninfected controls. Because EBV targets were undetectable in controls, Ct = 40, the final amplification cycle, was assigned to calculate the displayed fold-change estimates. (C) Immunoblots for EBNA1, Zta/BZLF1, EAD/BMRF1, VCA/p18, and GAPDH in uninfected, infected, and PMA/sodium-butyrate-treated infected cultures. Anti-IgG-induced lytic Akata cells were the positive control. GAPDH was the loading control. RT-qPCR data represent means ± SEM from ≥2 biological replicates, each assayed in technical triplicate. Statistical significance was assessed by *t* test versus the indicated control. *, *P* < 0.05; **, *P* < 0.01; ***, *P* < 0.001.

### RNA sequencing and preprocessing

RNA was isolated from FFPE tumor sections and cell cultures was isolated using Qiagen AllPrep (Cat# 80234) and assessed with an Agilent 2100 Bioanalyzer. GFP-enriched infected cultures and matched control cultures were collected five days after infection or mock exposure. Three independently established EBV-infected cultures and three independently maintained uninfected control cultures were used as biological replicates (n = 3 per group). Strand-specific poly(A)-selected libraries were prepared for the six cell-line samples and strand-specific rRNA-depleted libraries for the eight tumors. All were sequenced at the same facility on DNBSEQ with paired-end 100-base reads. FastQC 0.12.1 assessed read quality. fastp 1.1.0 performed adapter detection, six-base 5′ clipping, right-end trimming (four-base window; mean quality 20), and minimum-length filtering (35 bases) ^9^. MultiQC 1.35 summarized quality reports ^10^. Tumor analyses characterized viral sequences, whereas host differential-expression analysis used the cell-line libraries (Supplementary Methods S6).

### Human-Akata alignment and quantification

Reads were aligned to GRCh38 plus a coordinate-matched modified Akata EBV sequence (KC207813.1) ^11^ using STAR 2.7.10b ^12^, allowing up to 20 mapping loci and a mismatch-to-read-length ratio of 0.04. Genomic and transcriptome BAMs were generated. featureCounts 2.1.1 assigned host exon-level fragments to genes, requiring mapped mates and excluding chimeric, multimapping, and ambiguous assignments ^13^. Library orientation was assigned when one strand model yielded at least twice the alternative’s count. Otherwise, libraries were treated as unstranded. RSEM 1.3.3 estimated abundance from paired-end transcriptome alignments with the selected orientation and read-start-position-distribution estimation ^14^. EBV TPM values were summarized within library classes, and coverage was inspected in the Integrative Genomics Viewer (IGV) ^15^ (Supplementary Methods S7).

### Virome screening and normalization

A separate STAR alignment used GRCh38 plus 742 mammalian viral references, including Akata EBV type 1 and type 2 NC_009334.1 ^16^, allowing 50 loci and a mismatch ratio of 0.05. Per-reference screening excluded unmapped, secondary, supplementary, quality-control-failed, and duplicate-flagged records and required mapping quality ≥20. Species-aware EBV screening evaluated alternative alignments, counted accepted fragments once, and used mapping-quality thresholds of 1 (shared evidence) and 20 (strict evidence), aligned length ≥35 bases, and mismatch fraction ≤0.05. Shared fragments were not assigned to EBV type. Detailed filters are in Supplementary Methods S8.

Viral abundance was expressed as reads per million mapped human reads: RPMH = (qualifying viral read ends / mapped human read ends) × 10⁶, without a pseudocount. Viral numerators came from the broad virome alignment. Human denominators came from primary read-end alignments in the companion GRCh38-Akata BAM, excluding viral contigs and unmapped, secondary, supplementary, quality-control-failed, and duplicate-flagged records. Each mapped mate contributed one read. Unique and primary multimapping human alignments were included. Alternative EBV placements and same-genotype HPV references were not counted repeatedly. Library classes were summarized separately. Figure 3A displays log10(RPMH + 1), with 1 added only for visualization (Supplementary Methods S9).

**Figure 3.**
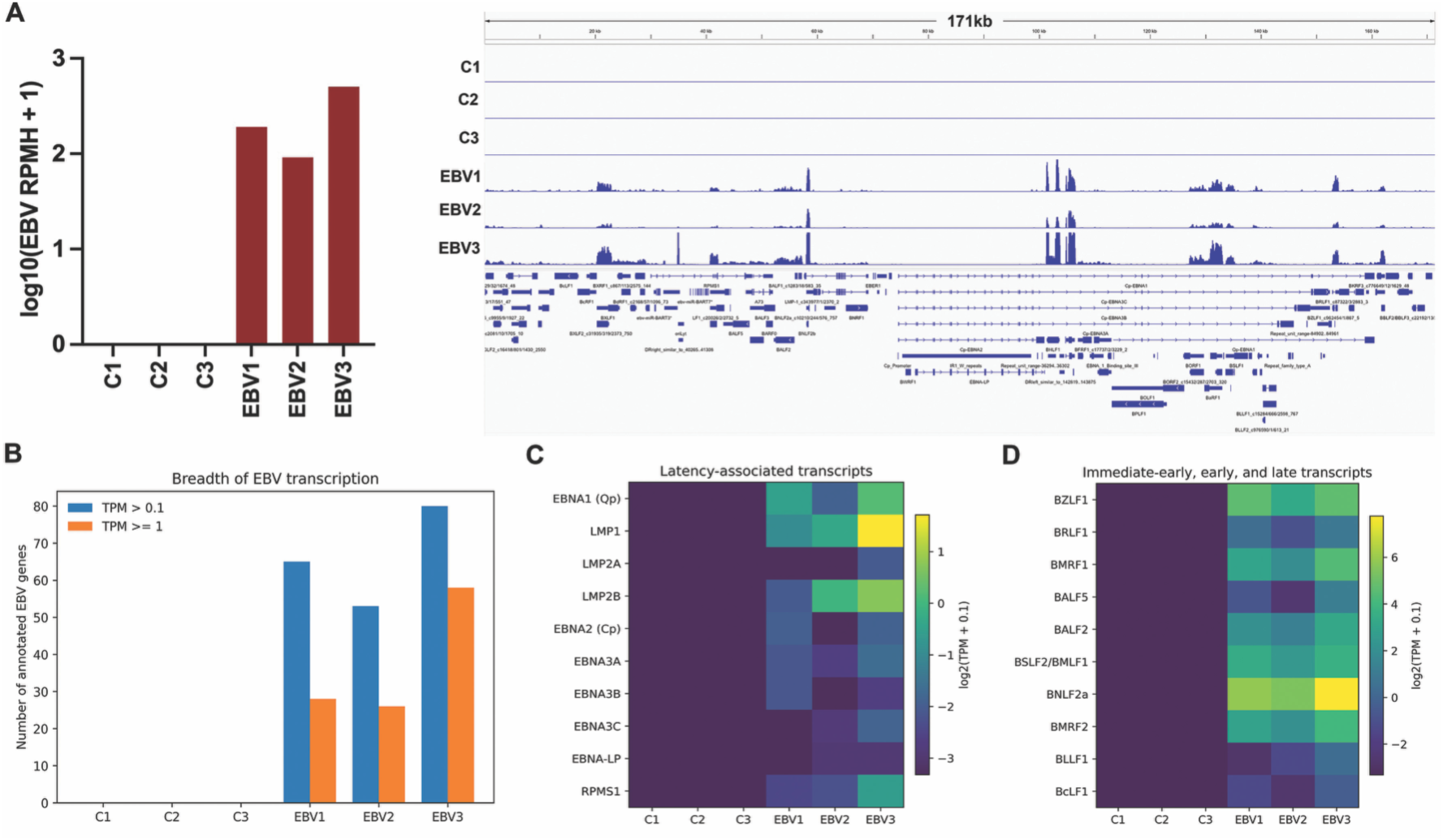
Latency-associated and lytic EBV transcription in infected conjunctival epithelial cultures. (A) EBV RNA-seq abundance in three independent control and three independently EBV-infected cultures (n=3 per group) (left), with representative read-coverage tracks across the modified Akata EBV reference (right). Bars display log10(RPMH + 1), where RPMH represents viral reads per million mapped human reads. RPMH values were calculated from qualifying viral and mapped human read-end counts without a pseudocount. The addition of 1 was applied only for logarithmic visualization. (B) Number of annotated EBV genes with TPM > 0.1 or TPM ≥ 1 in each culture. C1-C3 denote control cultures, and EBV1-EBV3 denote infected cultures. (C) Expression of selected latency-associated EBV transcripts and (D) immediate-early, early, and late lytic transcripts quantified using the modified Akata EBV reference. Heatmaps display log2(TPM + 0.1).

### Differential-expression and statistical analyses

DESeq2 compared three infected and three control cultures using integer featureCounts data, excluding EBV identifiers and retaining genes with ≥10 fragments in at least three libraries ^17^. The model was ∼ condition, with Wald tests and apeglm-shrunken display estimates ^18^. Wald statistics, P values, and adjusted P values came from unshrunk DESeq2 results. Benjamini-Hochberg FDR <0.05 defined significance ^19^. GSEA ranked eligible host genes by signed Wald statistic ^20^ and used fgseaMultilevel with MSigDB Hallmark sets of 15-500 genes ^21,22^.

Tumor EBER proportions were compared by two-sided Fisher exact test. Proportion intervals used Clopper-Pearson estimation, and odds ratios used Haldane-Anscombe correction. RT-qPCR t tests and replicate summaries are specified in the figure legends. RPMH values were descriptive and did not determine EBER classification. Analyses used R 4.4.2 and Python 3.11.15. Tests were two-sided unless specified otherwise (Supplementary Methods S10).

### Database accession number

The RNA-seq sequence data have been deposited in the NIH Sequence Read Archive (SRA) database under accession number PRJNA1544489.

## RESULTS

### Tumor-cell EBV detection is supported by complementary DNA and RNA assays

The clinical series comprised five Nigerian cSCC specimens from patients with HIV and three Japanese specimens from patients without HIV. Median ages were 30 years (range, 22-75) and 75 years (range, 56-93), respectively. Three Nigerian and all three Japanese patients were male (Figure 1A). HIV RT-qPCR results were concordant with the recorded patient classifications (Figure 1B).

EBER-ISH identified nuclear signals in epithelial tumor cells in four of five Nigerian specimens (cSCC-1, cSCC-3, cSCC-4, and cSCC-5), whereas the remaining Nigerian specimen and all three Japanese specimens were EBER-negative (Figure 1C). EBV DNA PCR detected viral DNA in cSCC-1, cSCC-4, and cSCC-5 (Figure 1D). After normalization to mapped human reads, EBV RNA-seq abundance was 2.68, 11.81, and 8.77 reads per million mapped human reads (RPMH) in cSCC-3, cSCC-4, and cSCC-5, respectively (Figure 1E). No EBV reads were detected in the remaining Nigerian specimens or in any Japanese tumor.

Each of the four EBER-positive tumors was supported by at least one additional nucleic-acid assay: cSCC-1 by DNA PCR, cSCC-3 by RNA-seq, and cSCC-4 and cSCC-5 by both DNA PCR and RNA-seq. The specimen-level assay matrix therefore connects spatial localization with complementary molecular evidence (Figure 1A). EBER positivity was 4/5 in Nigeria and 0/3 in Japan (two-sided Fisher exact P = 0.143).

Human papillomavirus (HPV) has also been investigated as a candidate contributor to ocular surface squamous neoplasia, with differing detection patterns across reported cohorts ^2,3^. Examining HPV alongside EBV can therefore provide additional virologic context for individual tumor series. HPV was assessed using targeted high-risk and combined high-/low-risk DNA PCR panels together with the broad RNA-seq virome screen. Neither PCR panel detected its target HPV genotypes in any of the eight tumors (Supplementary Figure S1B). Normalized RNA-seq abundance was 0.06 RPMH for HPV122 in Japanese specimen cSCC-6 and 0.08 RPMH for HPV41 in cSCC-7 (Supplementary Figure S1A). These assignments corresponded to six and eight qualifying read ends, respectively. HPV41 and HPV122 were not included in either PCR panel, so the negative targeted assays do not test these two sequencing assignments. These findings distinguish the panel-specific DNA results from the trace HPV-related RNA-seq evidence and provide complementary virologic context for the EBV-focused analysis.

### Establishment of an EBV-infected human conjunctival epithelial cell model

Primary conjunctival epithelial cells were immortalized by hTERT expression to generate HConEpiC-hTERT cells and then exposed to Akata-GFP-EBV. Infected cultures contained GFP-positive cells, providing a visual marker of infection (Figure 2A). GFP-based FACS was used to enrich the infected cell population for downstream analyses. RT-qPCR of sorted infected cells without PMA or sodium butyrate treatment detected EBER1, the latency-associated transcripts EBNA1 and LMP1, and the lytic transcripts BZLF1 and VCA/BcLF1, encoding Zta and the major capsid protein, respectively (Figure 2B).

Immunoblot analysis of untreated and PMA/sodium-butyrate-treated infected cultures further demonstrated viral protein expression, including EBNA1, Zta (BZLF1), EAD/BMRF1, and VCA/p18. Anti-IgG-induced lytic Akata cells served as a positive control. PMA/sodium-butyrate-treated infected cultures showed prominent Zta, BMRF1, and VCA/p18 signals, consistent with induction of the EBV lytic program (Figure 2C). Together, imaging, RNA assays, and protein detection established an experimentally accessible EBV-infected conjunctival epithelial system supporting viral gene expression and subsequent viral and host transcriptome profiling.

### EBV engages both latency-associated and lytic transcriptional programs

Virome screening distinguished the infected cultures from their controls. EBV RNA-seq abundance in infected cultures EBV1, EBV2, and EBV3 was 189.9, 90.3, and 505.4 RPMH, respectively, compared with 0 RPMH in controls C1, C2, and C3 (Figure 3A). Thus, the infection-associated contrast remained evident after normalization for mapped human read depth. Akata-EBV reference quantification identified 65, 53, and 80 annotated EBV genes above TPM 0.1 in the infected samples, including 28, 26, and 58 genes at TPM ≥ 1 (Figure 3B).

Latency-associated transcripts included Qp-EBNA1, LMP1, LMP2A/LMP2B, and RPMS1, together with low-level Cp-associated EBNA2/EBNA3 transcripts (Figure 3C). The viral profile also included BZLF1, BRLF1, BMRF1, BALF5, BALF2, BSLF2/BMLF1, BNLF2a, BMRF2, and late structural transcripts (Figure 3D). Across infected samples, BZLF1 abundance ranged from 9.77 to 25.78 TPM, BMRF1 from 3.60 to 20.94 TPM, and BNLF2a from 41.96 to 217.76 TPM. The coexistence of latency-associated and lytic transcripts at the culture level defines a mixed viral transcriptional state and provides a model in which both programs can be investigated.

### EBV infection induces inflammatory and epithelial-remodeling gene programs

Analysis of 17,607 filtered host genes identified 96 differentially expressed genes at FDR < 0.05, comprising 73 increased and 23 decreased transcripts in infected cultures. Sixty-two genes also had an absolute shrunken log2 fold change of at least 1 (Figure 4A-B). Prominent induced genes included ATF3 (shrunken log2 fold change [LFC], 3.65; FDR, 1.73 × 10⁻¹¹), NR4A2 (LFC, 3.55; FDR, 1.20 × 10⁻⁷), RGCC (LFC, 2.09; FDR, 1.20 × 10⁻⁷), and SERPINE2 (LFC, 1.30; FDR, 1.20 × 10⁻⁷). LOX (LFC, 1.89; FDR, 8.88 × 10⁻⁷), HES1 (LFC, 1.79; FDR, 3.43 × 10⁻⁶), DUSP1 (LFC, 2.36; FDR, 4.59 × 10⁻⁵), and ELF3 (LFC, 2.14; FDR, 4.59 × 10⁻⁵) were also increased. Induction of CXCL8 (LFC, 2.43; FDR, 0.00105) and CXCL2 (LFC, 2.59; FDR, 0.00251) identified a chemokine component of the epithelial response. Decreased transcripts included FGFBP1 (LFC, −2.05; FDR, 1.59 × 10⁻⁶) and CSTA (LFC, −1.35; FDR, 1.07 × 10⁻⁵). To validate the RNA-seq findings, RT-qPCR was performed for representative differentially expressed host genes (Figure 4C). Consistent with the RNA-seq results, HES1, CXCL8, CXCL2, and LOX were significantly upregulated following EBV infection, whereas FGFBP1, CSTA, SAPCD2, and TMEM40 were significantly downregulated. These RT-qPCR results provide orthogonal support for EBV-associated induction of inflammatory and epithelial-remodeling gene programs.

**Figure 4.**
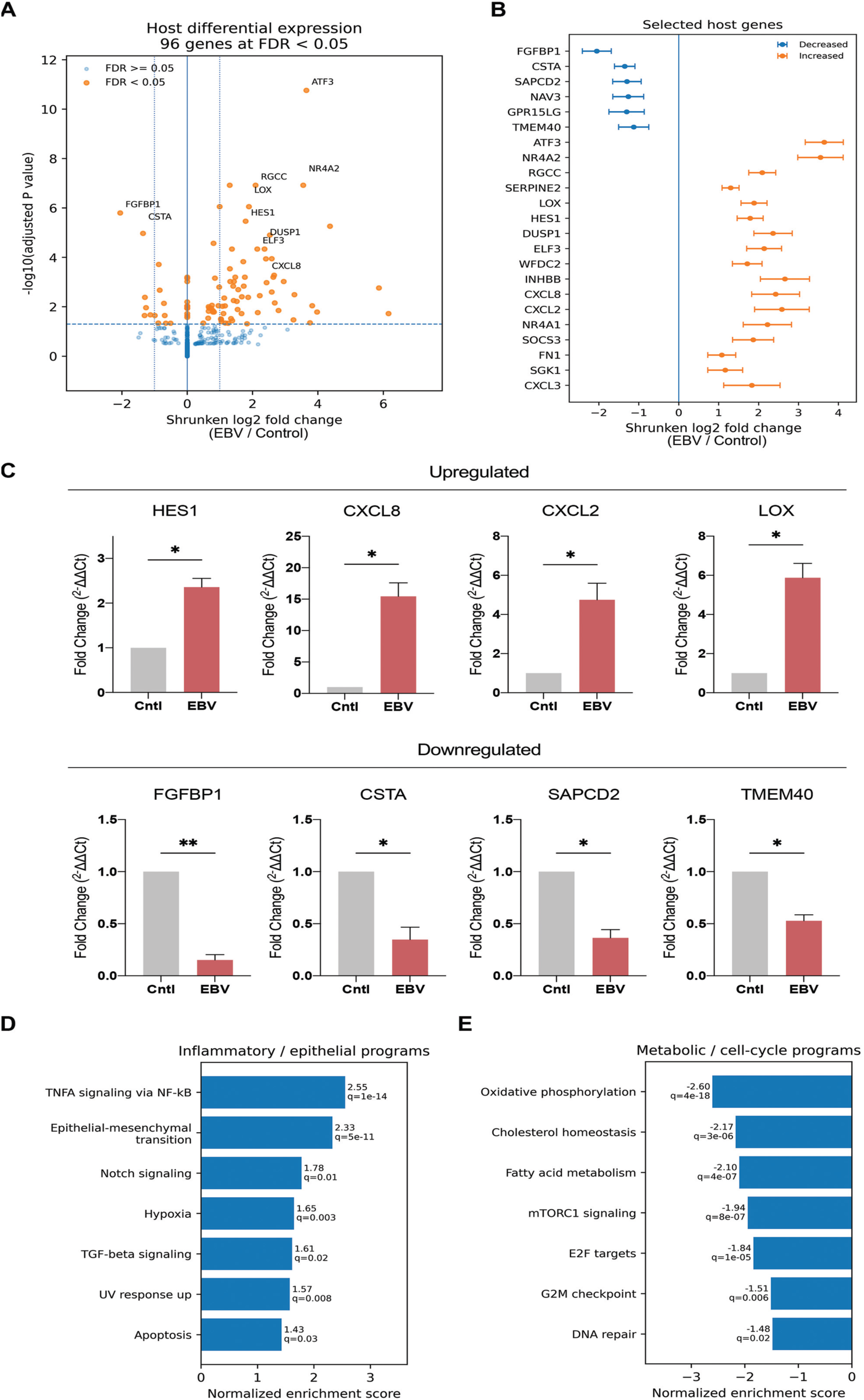
Host transcriptional reprogramming following in vitro EBV infection. (A) Volcano plot comparing three independently EBV-infected cultures and three independent uninfected control cultures (n = 3 biological replicates per group), showing shrunken log2 fold changes and -log10(adjusted P values). (B) Selected differentially expressed host genes. Error bars show the reported standard errors (lfcSE) of the shrunken estimates. (C) RT-qPCR validation of representative upregulated (HES1, CXCL8, CXCL2, and LOX) and downregulated (FGFBP1, CSTA, SAPCD2, and TMEM40) host genes. GAPDH was used as the internal reference gene, and relative expression was calculated using the 2^−ΔΔCt method, with uninfected control cells as the calibrator. (D) Positively enriched inflammatory and epithelial-state Hallmark gene sets and (E) negatively enriched metabolic and cell-cycle gene sets. Labels indicate normalized enrichment scores and false discovery rates (FDR). For RT-qPCR, data represent means ± SEM from ≥2 biological replicates; each biological replicate was assayed in technical triplicate. Statistical significance was assessed by *t* test versus the indicated control. *, *P* < 0.05; **, *P* < 0.01.

Hallmark analysis identified 23 enriched gene sets at FDR < 0.05 (Figure 4D-E). TNFA signaling via NF-κB (normalized enrichment score [NES], 2.55; FDR, 1.30 × 10⁻¹⁴) and epithelial-mesenchymal transition (NES, 2.33; FDR, 5.45 × 10⁻¹¹) showed prominent positive enrichment. Additional positively enriched sets included Notch signaling (NES, 1.78; FDR, 0.0130), hypoxia (NES, 1.65; FDR, 0.00270), TGF-β signaling (NES, 1.61; FDR, 0.0183), and UV response up (NES, 1.57; FDR, 0.00759).

This inflammatory and epithelial-state response was accompanied by coordinated metabolic and cell-cycle remodeling. Oxidative phosphorylation (NES, −2.60; FDR, 4.45 × 10⁻¹⁸), cholesterol homeostasis (NES, −2.17; FDR, 3.43 × 10⁻⁶), fatty-acid metabolism (NES, −2.10; FDR, 4.23 × 10⁻⁷), and mTORC1 signaling (NES, −1.94; FDR, 7.61 × 10⁻⁷) were negatively enriched. E2F targets, G2M checkpoint, and DNA-repair signatures also showed negative enrichment (Figure 4E). Together, the gene-level and pathway-level results identify a coordinated response to EBV involving inflammatory signaling, epithelial-state regulation, and metabolic adaptation.

## DISCUSSION

This study connects EBV detection in primary cSCC with in vitro infection and transcriptional reprogramming of human conjunctival epithelial cells. Its central contribution is the integration of tissue-localized viral evidence with a relevant epithelial model: EBER-positive tumor cells were identified in four HIV-associated Nigerian cSCC specimens, each with supporting viral DNA or RNA evidence, and infected conjunctival cultures displayed a defined host response. This approach advances the study of EBV in ocular surface cancer from viral detection toward experimentally testable mechanisms.

The tissue findings extend prior observations in African ocular surface neoplasia ^2–4^. The Zambian study detected EBV DNA and EBNA1 across preinvasive and invasive lesions, and the Zimbabwean case-control study identified an association between EBV DNA and cSCC ^2,4^. Tumor-cell EBER-ISH adds direct localization to epithelial tumor cells, while DNA PCR and RNA-seq provide complementary molecular evidence. The different contributions of these assays are informative: tissue imaging identifies the infected cellular compartment, PCR detects viral genomic material, and RNA-seq interrogates viral transcription. Expressing viral RNA abundance relative to mapped human reads adds a consistent sequencing-depth-adjusted measure to the spatial and DNA-based evidence. Their combined use strengthens the biological interpretation of EBV-associated tumor specimens without relying on any single detection platform.

HPV testing provides additional context for the EBV findings. Previous studies have evaluated HPV alongside EBV in ocular surface squamous neoplasia ^2,3^. In this series, targeted PCR detected none of the panel-covered genotypes, whereas RNA-seq identified trace HPV122- and HPV41-related sequences at 0.06 and 0.08 RPMH, respectively, in two Japanese tumors. Because these types were outside the PCR panels, the DNA and RNA findings address different genotype sets rather than conflicting assays of the same targets. The HPV sequence signals are therefore reported as secondary virome observations, while the central EBV evidence combines tumor-cell localization with an experimentally infected conjunctival epithelial model.

The clinical observations identify EBV in cSCC from patients with HIV, whereas the cell model defines epithelial responses to in vitro infection. These are complementary levels of evidence rather than interchangeable measures of tumor biology. The HIV-associated setting is particularly relevant in light of epidemiologic evidence connecting cSCC risk to immune dysfunction ^1^. These findings provide a basis for investigating how local epithelial infection, immune control, and environmental exposure may interact at the ocular surface.

Establishment of an EBV infection model derived from primary human conjunctival epithelial cells is a central advance of this study. To the authors’ knowledge, this is the first study to integrate experimental EBV infection with viral and host transcriptome profiling in hTERT-immortalized human conjunctival epithelial cells. Independently infected and control cultures permit these responses to be examined within a shared epithelial background. Normalized EBV RNA abundance ranged from 90.3 to 505.4 RPMH in infected cultures and 0 RPMH in controls, complementing the viral-gene and protein-expression findings. Published organotypic models have demonstrated the influence of epithelial organization and differentiation on EBV infection ^5^. The conjunctival system developed here extends this experimental framework to the ocular surface and enables focused investigations of infection-associated signaling, epithelial responses, and viral-state transitions.

A notable feature of this model is the engagement of both latency-associated and lytic programs. EBNA1 and LMP transcripts occurred alongside BZLF1, BRLF1, BMRF1, and a broader set of lytic transcripts. Protein detection and the response to PMA/sodium butyrate further support an active epithelial infection system. This configuration offers a means to investigate how latent and lytic components contribute to the host response. It is especially relevant to evidence that YAP/TAZ-associated epithelial signaling can regulate EBV reactivation through TEADs and that LMP1 can influence epithelial differentiation through related pathways ^6,7^.

Inflammatory cytokines provide another connection between host signaling and the EBV life cycle. A recent study from this group demonstrated that IL-17A directly triggers EBV lytic reactivation in latency I human B-cell models through IL-17RA-linked NF-κB/MAPK signaling ^23^. This finding, considered alongside the inflammatory transcriptional response observed here, motivates investigation of reciprocal interactions between inflammatory cues and EBV activity. The conjunctival epithelial model provides a system to define these interactions in an ocular surface context.

The host response places inflammatory signaling and epithelial remodeling at the center of the infection phenotype. TNFA/NF-κB and epithelial-mesenchymal-transition gene sets were among the strongest positive enrichments, accompanied by CXCL8 and CXCL2 induction and increased expression of LOX, SERPINE2, HES1, and immediate-response regulators such as ATF3 and NR4A2. This combination nominates a potential interaction between epithelial inflammatory output and regulation of cell state. The observed genes also provide practical candidates for testing changes in epithelial organization, matrix interactions, migration, and communication with neighboring cells.

TGF-β, Notch, hypoxia, and UV-response signatures further broaden the biological questions accessible in the model. Their enrichment suggests that EBV-responsive conjunctival cells may alter how they integrate differentiation signals and environmental stress. In the context of an exposed ocular surface, these responses raise the hypothesis that infection modifies epithelial adaptation to injury rather than acting solely through increased cell division. Experiments combining infection with ultraviolet exposure or defined inflammatory cues can directly test that hypothesis. Such studies would also clarify whether the nominated transcriptional programs influence survival, tissue repair, or persistent changes in epithelial behavior.

The accompanying decreases in oxidative phosphorylation, lipid-metabolic, and cell-cycle signatures are part of this coordinated reprogramming. They indicate that the infection-associated response includes redistribution of cellular activities, not simply induction of a single growth program. Determining how these metabolic and cell-cycle changes relate to viral state, infection duration, and longer-term epithelial behavior is an experimentally tractable next step. Time-resolved profiling, proliferation and clonogenic assays, and targeted perturbation of the leading host programs can distinguish adaptive responses from mechanisms that favor persistent epithelial dysfunction.

The integrated study design offers several strengths: spatial viral localization in tumors, complementary DNA and RNA assays, an epithelial model matched to the anatomic site, replicate-level infection comparisons, sequencing-depth-normalized viral RNA measurements, and parallel characterization of viral and host transcription. Together, these features prioritize a focused set of candidates for mechanistic analysis. Perturbing LMP1 or BZLF1, followed by measurement of NF-κB-associated transcription, chemokine output, and epithelial-state markers, would help assign viral contributions to the observed host programs. These experiments can be coupled to migration, clonogenicity, and three-dimensional epithelial assays to assess which responses are relevant to carcinogenesis.

In conclusion, EBV is localized to tumor cells in a subset of HIV-associated cSCC, and human conjunctival epithelial cells infected in vitro exhibit coordinated viral transcription and host inflammatory, epithelial-state, and metabolic responses. The study establishes a tissue-relevant platform for investigating EBV at the ocular surface and provides a molecular foundation for evaluating virus-host pathways with potential relevance to cSCC development and future intervention research.

## AUTHOR CONTRIBUTIONS

Conception and design: Hirotomo Dochi, Janardhan Avilala, and Zhen Lin. Development of methodology: Hirotomo Dochi, Janardhan Avilala, and Zhen Lin. Acquisition of data: Hirotomo Dochi, Janardhan Avilala, Maria Morales, and Zhen Lin. Analysis and interpretation of data (e.g., statistical analysis, biostatistics, computational analysis): Hirotomo Dochi, Janardhan Avilala, and Zhen Lin. Writing, review, and/or revision of the manuscript: Hirotomo Dochi, Janardhan Avilala, Ramsy Abdelghani, David Becnel, Mark Sides, Gilbert F. Morris, Wenke Feng, Hong Liu, and Zhen Lin. Administrative, technical, or material support (i.e., reporting or organizing data, constructing databases): Yutaro Yamada, Masayuki Takahira, Satoru Kondo, Tomokazu Yoshizaki, and Zhen Lin. Study supervision: Zhen Lin

## ACKNOWLEDGEMENTS

Research reported in this publication was supported by a U.S.-Japan Cooperative Medical Sciences Program Collaborative Award from the National Institute of Allergy and Infectious Diseases and CRDF Global (grant number DAA3-19-65602-1), a Louisiana Cancer Research Center seed grant, a Carol Lavin Bernick faculty grant, and a Ladies Leukemia League research grant to ZL.

## CONFLICT OF INTEREST

The authors declare no conflicts of interest.

## DATA AVAILABILITY STATEMENT

All the data shown in this article are available from the corresponding author upon reasonable request.

## USE OF ARTIFICIAL INTELLIGENCE-ASSISTED TOOLS

ChatGPT (OpenAI) was used solely to assist with grammar, language, and readability. All scientific content was developed, reviewed, and approved by the authors.

---

**Supplementary Figure S1.**
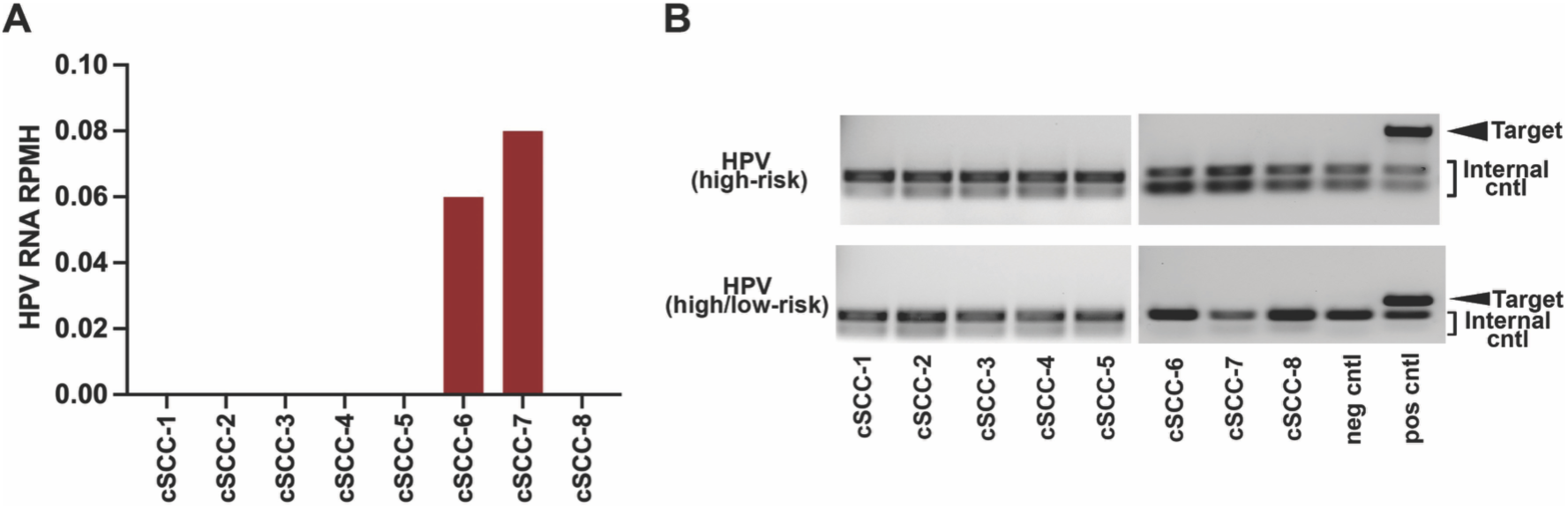
Targeted HPV DNA testing and HPV-related RNA-seq evidence. (A) Genotype-specific HPV RNA-seq abundance expressed as viral reads per million mapped human reads (RPMH) on a linear scale. HPV122 abundance was 0.06 RPMH in cSCC-6, and HPV41 abundance was 0.08 RPMH in cSCC-7. HPV41/HPV122 reads were not detected in the other tumors. (B) HPV DNA PCR using the high-risk and combined high-/low-risk panels, with the indicated positive controls. None of the tested tumors was positive for the panel-covered genotypes. HPV41 and HPV122 were outside the scope of both panels, so the negative PCR results do not assess these sequencing assignments. Cases cSCC-1 to cSCC-5 are from Nigeria and cSCC-6 to cSCC-8 are from Japan. The RNA-seq findings are described as trace HPV-related sequence evidence.

## Supplementary Methods

### S1. EBER in situ hybridization

Five-micrometer FFPE sections were prepared for hematoxylin and eosin staining and Epstein-Barr virus (EBV)-encoded RNA in situ hybridization (EBER-ISH). EBER-ISH was performed using the HistoSonda EBER XISH Probes kit (American MasterTech). Briefly, the FFPE tissue sections were deparaffinized, rehydrated in a graded solution of xylene and alcohol, and deproteinized with proteinase K. Samples were incubated with a digoxigenin EBER probe and washed with deionized water and 1× PBS. They were incubated first with anti-digoxin antibody and horseradish peroxidase-conjugated anti-mouse antibody and subsequently with 3,3′-diaminobenzidine (Biocare Medical, Cat# BDB2004), counterstained with hematoxylin (StatLab, Cat# SL95-1), and washed again with 1× PBS. Slides were then dehydrated in a graded solution of xylene and alcohol and subsequently sealed with the VectaMount permanent mounting medium (Vector Laboratories, Cat# H-5000). An EBV-positive and an EBV-negative control were included in each run. Digital slide scanning was performed with a Motic EasyScan Pro 6 digital slide scanner (Motic, Hong Kong, China), and images were captured at 40× magnification. EBER positivity was defined as discrete nuclear signal in morphologically identifiable epithelial tumor cells. Tumor-infiltrating lymphocytes were evaluated separately to distinguish tumor-cell-associated EBV from signal in bystander immune cells. EBER-ISH was selected as the primary variable for assigning tumors to EBV-positive and EBV-negative groups because it provides spatial localization of EBV in tissue (8).

### S2. HIV, EBV, and HPV nucleic-acid assays

DNA and RNA were extracted from FFPE tumor sections using the Qiagen AllPrep kit according to the manufacturer’s instructions. HIV status was established clinically and was additionally evaluated in tumor-derived RNA using the HIV TaqMan RT-PCR kit (Norgen Biotek, Cat# TM33750). Real-time PCR was carried out following the manufacturer’s instructions. Fluorescent signals were detected in the FAM (target) and HEX (internal control) channels. The thermal cycling conditions were as follows: 50 °C for 30 min, 95 °C for 3 min, followed by 40 cycles of 95 °C for 15 s and 60 °C for 30 s.

EBV genomic DNA was detected by qPCR targeting the BALF5 locus. qPCR reactions were performed using a 2x SYBR Green Master Mix (Bio-Rad) on a Bio-Rad real-time PCR system. The thermal cycling conditions were as follows: 95 °C for 5 min, followed by 40 cycles of 95 °C for 15 s and 60 °C for 1 min. Amplicons were visualized by gel electrophoresis. Primer sequences are provided in Supplementary Table S1. Human genomic tubulin served as an extraction and amplification control.

HPV DNA was evaluated using two commercial multiplex endpoint PCR assays (Norgen Biotek): a high-risk HPV panel targeting HPV genotypes 16, 18, 31, 33, 35, 39, 45, 51, 52, 56, 58, 59, and 68 (Cat# EP32200), and a combined high- and low-risk HPV panel (Cat# EP31500). The latter detects 13 high-risk HPV genotypes (HPV16, 18, 31, 33, 35, 39, 45, 51, 52, 56, 58, 59, and 68) and five low-risk genotypes (HPV6, 11, 42, 43, and 44). Together, these assays cover 18 distinct HPV genotypes. PCR amplification was performed according to the manufacturer’s instructions, with HPV-positive control DNA, no-template controls, and an internal human DNA control included to assess assay performance. PCR products were separated by agarose gel electrophoresis, and HPV detection was determined by the presence of amplification products of the expected sizes. HPV41 and HPV122, the two genotypes identified at low abundance by RNA-seq, were not included in either PCR panel. Therefore, negative PCR results indicated that the targeted HPV genotypes were not detected under the assay conditions and did not exclude infection with HPV genotypes outside the panels’ coverage.

### S3. Conjunctival epithelial cells and EBV infection

Primary human conjunctival epithelial cells (HConEpiC, ScienCell, Cat# 6630) were cultured in corneal epithelial cell medium at 37 °C under humid conditions and immortalized by lentiviral delivery of human telomerase reverse transcriptase (hTERT). The resulting HConEpiC-hTERT line was selected with hygromycin.

Immortalized cells were then infected with cell-free Akata-GFP-EBV (6.7 × 10⁵ Raji IU/mL) at a multiplicity of infection (MOI) of 0.17, by adding 100 µL of viral supernatant to 2 mL of culture medium. Mock-infected cultures underwent the same handling without infectious virus. Three independently EBV-infected cultures and three independently maintained uninfected control cultures were used as biological replicates (n = 3 per group) and harvested five days after infection or mock exposure. The infected cell population was enriched by fluorescence-activated cell sorting (FACS) based on GFP expression. For lytic reactivation experiments, EBV-infected cells were treated with phorbol 12-myristate 13-acetate (PMA; 20 ng/mL; Sigma-Aldrich, Cat# P1585) and sodium butyrate (SB; 3 mM; Sigma-Aldrich, Cat# B5887) for 48 hours before harvest.

### S4. RT-qPCR

cDNA was synthesized from total RNA using iScript gDNA Clear cDNA Synthesis Kit (Bio-Rad, Cat# 172-5035). qPCR analysis was performed using iQ SYBR Green Supermix (Bio-Rad, Cat# 170-8882) on a Bio-Rad CFX96 instrument as follows: 1 μl of cDNA and 1 μl of 10 μM primers were mixed with 10 μl of SYBR green supermix and 8 μl nuclease-free H2O to a 20 μl reaction volume. Polymerase was activated and cDNA was denatured at 95°C for 5 minutes. cDNA was then amplified for 40 cycles with 15 s denaturation at 95°C, 60 s annealing/extension and plate reading at 60°C. Melting curve analysis was performed at temperatures from 60°C to 90°C with 0.5°C increment per 5 s. Technical triplicates were averaged for each biological sample before calculating group means. Primer sequences are listed in Supplementary Table S1.

GAPDH served as the internal reference gene, and relative transcript expression was calculated using the 2^−ΔΔCt^ method with uninfected controls as the calibrator. For Figure 2B, the viral targets were EBER1, BZLF1, EBNA1, LMP1, and VCA/BcLF1. Infected cells were GFP-positive after FACS and had not received PMA or sodium butyrate. EBV targets were not detected in uninfected controls. Ct = 40, the final amplification cycle, was assigned to calculate the displayed fold-change estimates. Figure 4C used GAPDH normalization to assess the indicated host transcripts.

### S5. Immunoblotting

The cells were pelleted and washed with 1X PBS and then the cells were immediately suspended in 100 μl of 1× Laemmli sample buffer and heated for 15 minutes at 95°C. Protein concentrations of the whole-cell extracts were measured with the Bio-Rad protein assay kit according to the manufacturer’s instructions. Equal amounts of protein were subjected to SDS-polyacrylamide gel electrophoresis and transferred to PVDF membranes (Whatman). The blots were blocked for 60 minutes in Tris-buffered saline containing 5% nonfat powdered milk and 1% FBS and then incubated with the primary antibody (in blocking buffer) overnight at 4°C. The blots were washed three times with 1× TBST (140 mM NaCl, 3 mM KCl, 25 mM Tris-HCl [pH 7.4], 0.1% Tween 20) (each wash was carried out for approximately 15 minutes). To detect the chemiluminescence signals, the blots were then incubated with horseradish peroxidase-conjugated secondary antibody (Bio-Rad) in blocking buffer for 1 h at room temperature. Blots were washed as described above and analyzed with an enhanced chemiluminescence detection system (Perkin-Elmer) according to the manufacturer’s recommendations. Signals were acquired using a Bio-Rad ChemiDoc imaging system or by exposure to Fuji Super RX films (Fujifilm) followed by film scanning. To detect infrared fluorescence signals, an Odyssey infrared imaging system (Li-Cor Biosciences) was used with a secondary antibody (IRDye 800CW goat anti-mouse antibody; Li-Cor, Cat# 926-32210). Immunoblot figure panels were assembled from the original image files using uniform brightness/contrast or levels adjustments applied to the entire blot image or panel. No selective enhancement, erasure, cloning, masking, background filling, or local manipulation was performed.

The following primary antibodies were used for Western blot analysis: anti-Zta MAb (Santa Cruz, Cat# sc-53904), anti-BMRF1 MAb (Santa Cruz, Cat# SC-58121), anti-VCA/p18 (ThermoFisher, Cat# PA1-73003), anti-EBNA1 (Santa Cruz, Cat# sc-81581) and anti-GAPDH (Santa Cruz, Cat# SC-47724).

### S6. RNA library preparation and preprocessing

Total RNA was isolated from FFPE tumor sections and cell cultures using the Qiagen AllPrep kit (Qiagen, Cat# 80234). GFP-enriched infected cultures and matched uninfected controls were collected five days after infection or mock exposure. Three independently established EBV-infected cultures and three independently maintained uninfected control cultures were used as biological replicates (n = 3 per group). RNA quality was assessed with an Agilent 2100 Bioanalyzer. Strand-specific poly(A)-selected libraries were prepared for the six cell-line samples, and strand-specific rRNA-depleted libraries were prepared for the eight tumor specimens. Libraries were sequenced at the same facility on a DNBSEQ platform to generate paired-end 100-base reads.

Raw-read quality was assessed with FastQC (version 0.12.1). Reads were processed with fastp (version 1.1.0), including paired-end adapter detection, removal of the first six bases from each mate, right-end quality trimming with a four-base window and mean-quality threshold of 20, and a minimum retained read length of 35 bases (9). Quality reports were summarized with MultiQC (version 1.35) (10). Tumor RNA-seq analyses reported here characterized viral sequences. Host differential-expression analysis used the six cell-line libraries.

### S7. Human-Akata EBV alignment and expression quantification

Reads were aligned to a composite reference comprising the GRCh38 primary human assembly and a coordinate-matched modified Akata EBV sequence derived from accession KC207813.1 (11). STAR version 2.7.10b was used with a maximum of 20 mapping loci and a maximum mismatch-to-read-length ratio of 0.04 (12). Coordinate-sorted genomic BAM files and transcriptome-coordinate alignments were generated with TranscriptomeSAM and GeneCounts output and the NH, HI, AS, nM, and MD alignment attributes.

Host exon-level fragments were assigned to gene identifiers using featureCounts version 2.1.1 (13). Paired fragments were counted with both mates mapped, chimeric pairs excluded, and default exclusion of multimapping and ambiguously overlapping assignments. Library orientation was evaluated by comparing forward- and reverse-stranded counts. An orientation was selected when its assigned count was at least twice that of the alternative, otherwise the library was treated as unstranded. RSEM version 1.3.3 estimated human and EBV abundance from paired-end transcriptome alignments with the selected strandedness and read-start-position-distribution estimation (14). EBV transcripts per million (TPM) were summarized within each library class. Viral genomic coverage was inspected using the Integrative Genomics Viewer (15).

### S8. Broad virome screening and EBV-specific evidence

A separate composite reference combined GRCh38 with a curated panel containing 742 mammalian viral reference sequences, including Akata EBV type 1 and EBV type 2 accession NC_009334.1 (16). STAR alignment allowed up to 50 mapping loci and a maximum mismatch-to-read-length ratio of 0.05. Viral-contig alignments were extracted for read-level analysis. Per-reference summaries excluded unmapped, secondary, supplementary, quality-control-failed, and duplicate-flagged alignments and required mapping quality of at least 20. For fragment-level screening summaries, paired reads were counted once by fragment identifier. Covered positions, genome breadth, and coverage across 500-base bins were recorded. Read-end counts were then used for the normalized abundance measure described below.

To accommodate conserved reads shared between EBV type 1 and type 2 references, a species-aware analysis evaluated reported alternative alignments and counted each accepted fragment once at the EBV-species level. This analysis used a relaxed mapping-quality threshold of 1 for shared EBV evidence, a threshold of 20 for strict evidence, a minimum aligned length of 35 bases, and a maximum mismatch fraction of 0.05. The NH attribute was used to identify unobserved alternative alignments. Species-shared fragments were not interpreted as type-specific evidence. RNA-seq results were assessed alongside tumor-cell EBER-ISH, EBV DNA PCR, and the Akata-EBV reference coverage review.

### S9. Normalization of viral RNA-seq abundance

Viral RNA-seq abundance was expressed as viral reads per million mapped human reads (RPMH). For each sample, the human denominator was obtained from primary read-end alignments to human reference sequences in the companion GRCh38-Akata genomic BAM. Human reference sequences were defined using the human-only FASTA index, excluding viral contigs. Unmapped, secondary, supplementary, quality-control-failed, and previously duplicate-flagged records were excluded. Both uniquely mapped human reads and the primary representation of multimapping human reads were included. Each mapped mate contributed one read. The denominator was therefore a genomic alignment count rather than a gene-assigned fragment count.

EBV numerators comprised distinct qualifying read ends within fragments accepted by the species-aware type 1/type 2 virome screen. Alternative placements of an accepted read end on the two EBV references were not counted repeatedly. HPV numerators comprised qualifying primary viral read ends with mapping quality of at least 20, with read identifiers combined without duplication when multiple references represented the same HPV genotype. Viral numerators came from the broad virome alignment, whereas all human denominators came consistently from the companion GRCh38-Akata alignment. Normalized abundance was calculated as: RPMH = (qualifying viral read ends / mapped human read ends) × 10⁶. Normalization was performed without adding a pseudocount. Cell-line and tumor abundances were summarized separately within their respective library classes. For Figure 3A, bars display log10(RPMH + 1). The addition of 1 was applied only for visualization. RPMH was distinct from RSEM gene-level TPM and did not replace the integer host-gene counts used for DESeq2.

### S10. Differential-expression and statistical analyses

Host differential expression was analyzed using integer featureCounts data with EBV identifiers removed. Genes with at least 10 fragments in three or more cell-line libraries were retained. DESeq2 modeled the infected-versus-control comparison using the formula ∼ condition and Wald tests (17). For displayed effect sizes, log2 fold changes were shrunk using apeglm (18). Wald statistics, P values, and adjusted P values were taken from the unshrunk DESeq2 analysis. The Benjamini-Hochberg method controlled the false discovery rate (FDR), with FDR < 0.05 defining statistical significance (19). Gene-set enrichment analysis (GSEA) used the signed DESeq2 Wald statistic to rank the full eligible host-gene list (20). Hallmark sets from MSigDB were analyzed using fgseaMultilevel, retaining sets of 15-500 genes (21, 22).

Categorical comparisons of tumor EBV prevalence used a two-sided Fisher exact test. Exact two-sided Clopper-Pearson confidence intervals were calculated for cohort-specific proportions. Because one cell of the 2 by 2 table was zero, the odds ratio and confidence interval were estimated using the Haldane-Anscombe correction. All transcriptomic P values were adjusted for multiple testing by the Benjamini-Hochberg method (19). Viral RPMH values were reported descriptively for individual samples. Normalization did not alter the underlying viral-evidence assignments or the tumor EBER-ISH comparison. Analyses were performed with R (version 4.4.2) and Python (version 3.11.15). All statistical tests were two-sided unless otherwise specified.

**Supplementary Table S1:**
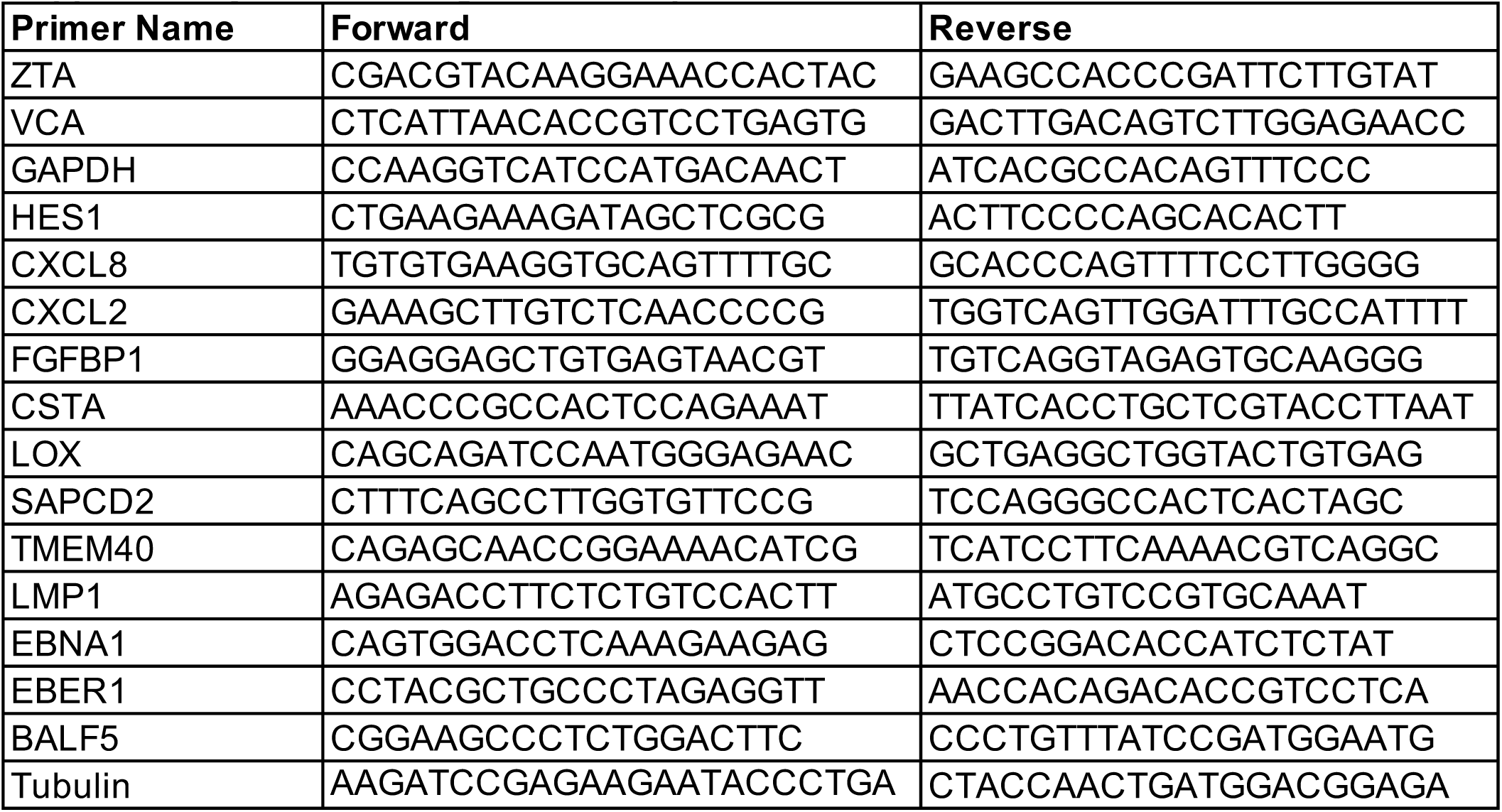
Oligonucleotide primers used.

## Notes

### Competing Interest Statement

The authors have declared no competing interest.

